# Membrane voltage-dependent activation of the flagellar protein export engine

**DOI:** 10.1101/2020.07.18.210377

**Authors:** Tohru Minamino, Yusuke V. Morimoto, Miki Kinoshita, Keiichi Namba

## Abstract

Ion motive force (IMF) consists of the electric potential difference (ΔΨ) and the ion concentration difference (ΔpI) across the cytoplasmic membrane. The flagellar protein export machinery is an ion/protein antiporter utilizing IMF to drive ion-coupled protein export, but it remains unknown how. Here, we report a ΔΨ-dependent activation mechanism of the transmembrane export gate complex. Depletions of both H^+^ and Na^+^ gradients nearly diminished flagellar protein export in the absence of the cytoplasmic ATPase complex, but an increase in ΔΨ by an upward shift of external pH from 7.5 to 8.5 dramatically recovered it. An increase in the cytoplasmic level of export substrates and gain-of-function mutations in FlhA enhanced protein export at external pH 7.5 in the absence of Na^+^ in a similar manner to ΔΨ increase. We propose that the export gate complex has a voltage-gated mechanism to activate the ion/protein antiporter of the flagellar protein export engine.

---

Ion motive force (IMF) across the biological membrane is one of the most important biological energies. IMF is composed of the electric potential difference (ΔΨ) and chemical potential difference of ions (ΔpI) across the membrane. IMF is utilized for many of the essential biological activities, such as ATP synthesis, solute transport, nutrient uptake, protein secretion, flagella-driven motility and so on^1^. Although ΔΨ and ΔpI are equivalent driving forces for the translocation of ions across the cytoplasmic membrane, it has been reported that ΔΨ facilitates the translocation of negatively charged residues of secreted proteins across the membrane by an electrophoretic mechanism^2,3^. Thus, ΔΨ plays a distinct role in protein secretion.

The bacterial flagellum is a supramolecular protein complex consisting of the basal body acting as an ion-driven rotary motor, the hook as a universal joint and the filament as a helical propeller. The flagellar motor converts the ion influx through a transmembrane ion channel of the stator unit into the force for high-speed rotation of the long helical filament^4,5^. It has been reported that ΔΨ and ΔpI are also not equivalent as the driving force for high-speed motor rotation at low load^6^.

For construction of the hook and filament structures in the cell exterior, flagellar building blocks are transported via a specialized protein export apparatus to the distal end of the growing flagellar structure. The flagellar protein export machinery of *Salmonella enterica* serovar Typhimurium (hereafter referred to *Salmonella*) is composed of a transmembrane export gate complex made of FlhA, FlhB, FliP, FliQ and FliR and a cytoplasmic ATPase ring complex consisting of FliH, FliI and FliJ (Fig. 1)^7,8^. These proteins are evolutionally related to those of virulence-associated type III secretion systems of pathogenic bacteria, which inject effector proteins into eukaryotic host cells for invasion^9^. Furthermore, the entire structure of the cytoplasmic ATPase ring complex is structurally similar to a cytoplasmic F_1_ part of F_O_F_1_-ATPsynthase, which utilizes PMF for ATP synthesis^10,11^.

**Fig. 1.**
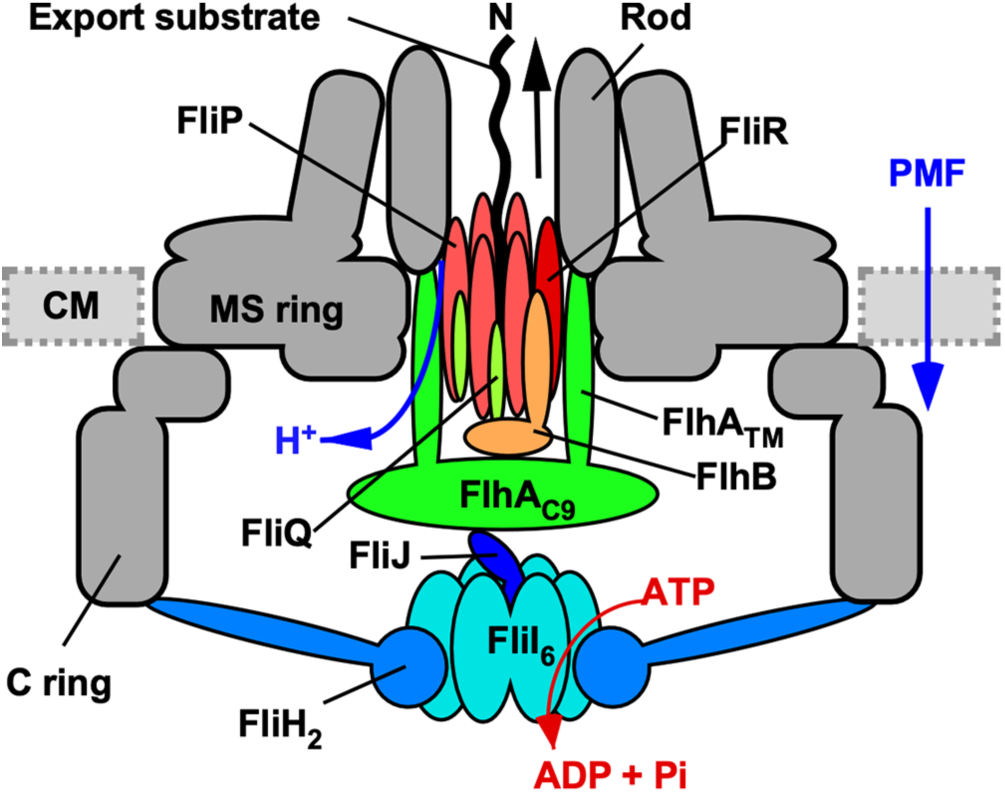
Cartoon of the flagellar protein export machinery. The flagellar protein export machinery is composed of a transmembrane export gate complex made of FlhA, FlhB, FliP, FliQ and FliR and a cytoplasmic ATPase ring complex consisting of FliH, FliI and FliJ. The export gate complex is located inside the MS ring and utilizes proton motive force (PMF) across the cytoplasmic membrane (CM) to drive proton (H^+^)-coupled flagellar protein export. FliP, FliQ and FliR form a polypeptide channel complex. FlhB associates with the FliP/FliQ/FliR complex, and its C-terminal cytoplasmic domain (FlhB_C_) projects into the central cavity of the C ring. FlhA forms a homo-nonamer through interactions between the C-terminal cytoplasmic domain of FlhA (FlhA_C_), and its N-terminal transmembrane domain (FlhA™) forms a pathway for the transit of H^+^ from the periplasm to the cytoplasm. The cytoplasmic ATPase ring complex associates with the C ring through an interaction between FliH and a C ring protein, FliN. ATP hydrolysis by the FliI ATPase activates the export gate complex through an interaction between FliJ and FlhA_C_, allowing the gate complex to become an active export engine to couple the proton flow through the FlhA proton channel to the translocation of export substrates into the polypeptide channel. When the cytoplasmic ATPase complex is missing, the export gate complex also utilizes Na^+^ as the coupling ion to drive flagellar protein export.

The flagellar protein export machinery utilizes ATP and PMF to drive flagellar protein export^12,13^. The transmembrane export gate complex couples an inward-directed proton (H^+^) flow with an outward-directed protein export^14,15^. ATP hydrolysis by the FliI ATPase activates the export gate complex to become an active export engine that uses PMF to drive H^+^-coupled protein export^16^. This export engine can also use sodium (Na^+^) motive force (SMF) to drive Na^+^-coupled flagellar protein export over a wide range of external pH when the cytoplasmic ATPase complex does not function properly^17^. It has been shown that FlhA forms a pathway for the transit of both H^+^ and Na^+^ across the cytoplasmic membrane^17^, suggesting that FlhA acts as an energy supplier for the export gate complex.

Only the ΔΨ component of PMF is sufficient for flagellar protein export by *Salmonella* wild-type cells^13,14^. However, a chemical potential gradient of either H^+^ (ΔpH) or Na^+^ (ΔpNa) becomes essential in the absence of the cytoplasmic ATPase complex, suggesting that ΔΨ and ΔpH/ΔpNa are used for different steps of the flagellar protein export process^14,17^. To clarify how the export gate complex uses these two distinct energies, we used a *Salmonella* Δ*fliH-fliI flhB(P28T)* mutant, of which flagellar protein export engine requires both ΔΨ and ΔpH/ΔpNa to exert its protein transport activity^14,17^. We show that an increase in ΔΨ by an upward shift of external pH from 7.5 to 8.5 activates flagellar protein export by this mutant in the absence of ΔpH and ΔpNa, suggesting the presence of a ΔΨ-dependent activation mechanism of the export gate complex to transport flagellar building blocks to form flagella on the cell surface.

## Results

### Effect of increase in ΔΨ on flagellar protein export

The ΔpH and ΔpNa components are thought to be required for efficient transit of H^+^ and Na^+^ through the FlhA ion channel, respectively, besides **ΔΨ** when FliH and FliI are missing^14,17^. In *Salmonella*, intracellular pH is maintained at about 7.5 over a wide range of external pH^18^. Higher external pH than 7.5 results in a negative ΔpH but total PMF is maintained by an increase in ΔΨ in bacterial cells^19,20^. Consistently, our measurements showed that ΔΨ was larger at external pH 8.5 than that at external pH 7.5 (Fig. 2a and Supplementary Table 1) and that there was no significant difference in total PMF (Fig. 2b and Supplementary Table 1). To clarify the role of ΔΨ in flagellar protein export, we used a *Salmonella* Δ*fliH-fliI flhB(P28T)* strain, of which *flhB(P28T)* mutation considerably increases the protein transport activity of the transmembrane export gate complex in the absence of FliH and FliI^12^. The Δ*fliH-fliI flhB(P28T)* mutant cells were exponentially grown in T-broth at external pH 7.5 (TB-7.5) or 8.5 (TB-8.5) in the absence of NaCl, and then the levels of the hook-capping protein FlgD secreted by this mutant were analyzed by immunoblotting with polyclonal anti-FlgD antibody (Fig. 3a) as a representative assay of flagellar protein export activity. In the Δ*fliH-fliI flhB(P28T)* Δ*flhA* strain, which is a negative control, no FlgD was detected in the culture supernatant. The Δ*fliH-fliI flhB(P28T)* mutant cells secreted FlgD into the culture media at external pH 8.5 but not at external pH 7.5 (Fig. 3a), indicating that about 1.5-fold greater ΔΨ activates flagellar protein export in the absence of the cytoplasmic ATPase complex. When the external pH value was varied over a range of 7.0 to 8.5, the relative secretion level of FlgD by this mutant increased from almost zero at pH 7.0 and 7.5 to about 0.8 at external pH 8.0 and finally to 1.0 at external pH 8.5 (Fig. 3b). Because there was no difference in total PMF between external pH values of 7.5 and 8.5 (Fig. 2b), we suggest that the export gate complex can become an active ion/protein antiporter to drive flagellar protein export in a ΔΨ-dependent manner when ΔΨ is increased to a level above a certain threshold.

**Fig. 2.**
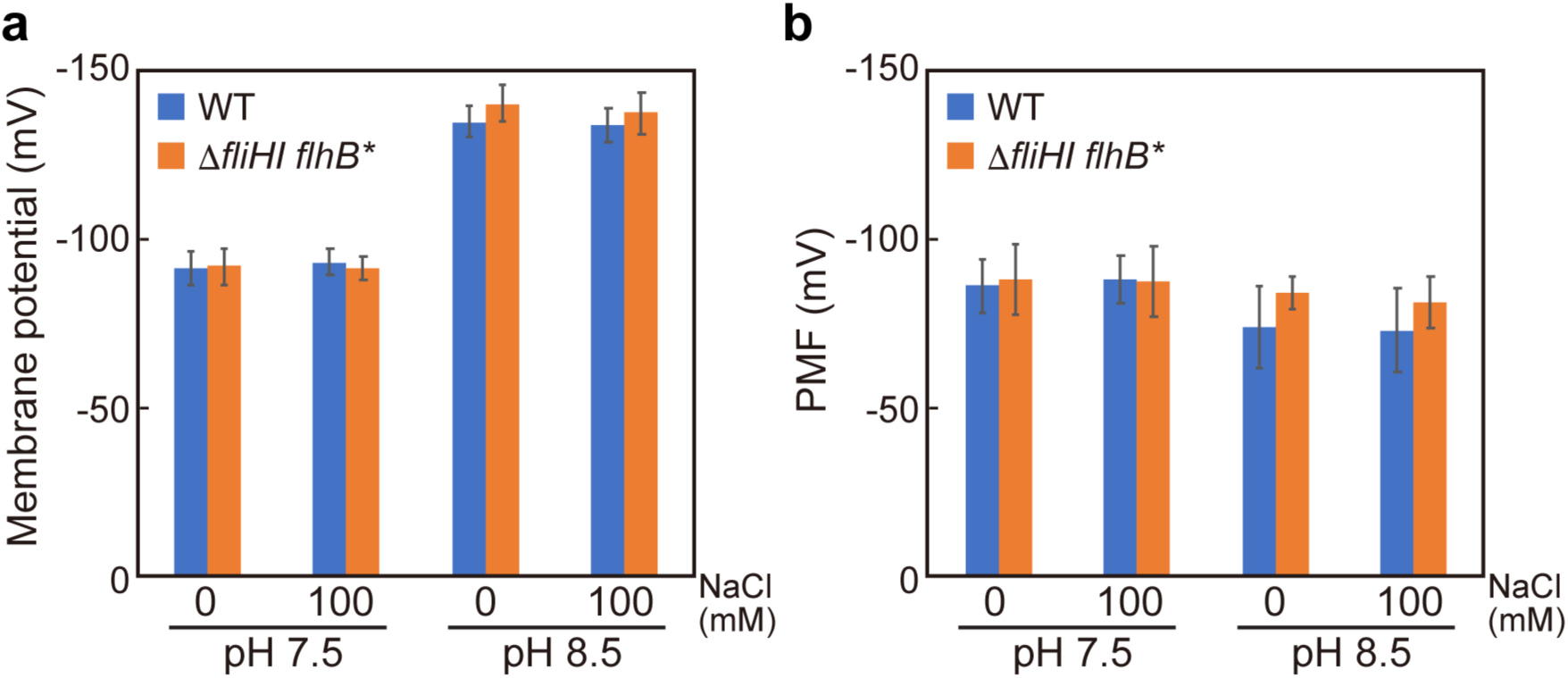
Effect of external pH on ΔΨ. **(a)** Measurements of ΔΨ. *Salmonella* SJW1103 (WT) and MMHI0117 (Δ*fliHI flhB*\*) cells were exponentially grown at 30°C in TB-7.5 or TB-8.5 with or without 100 mM NaCl. The membrane potential difference (mV) was measured by using tetramethylrhodamine methyl ester. Vertical bars indicate standard deviations. **(b)** Measurements of total proton motive force (PMF). Intracellular pH was measured using pHluorin(M153R). Four independent measurements were carried out. Vertical bars indicate standard deviations. (See Supplementary Table 1)

**Fig. 3.**
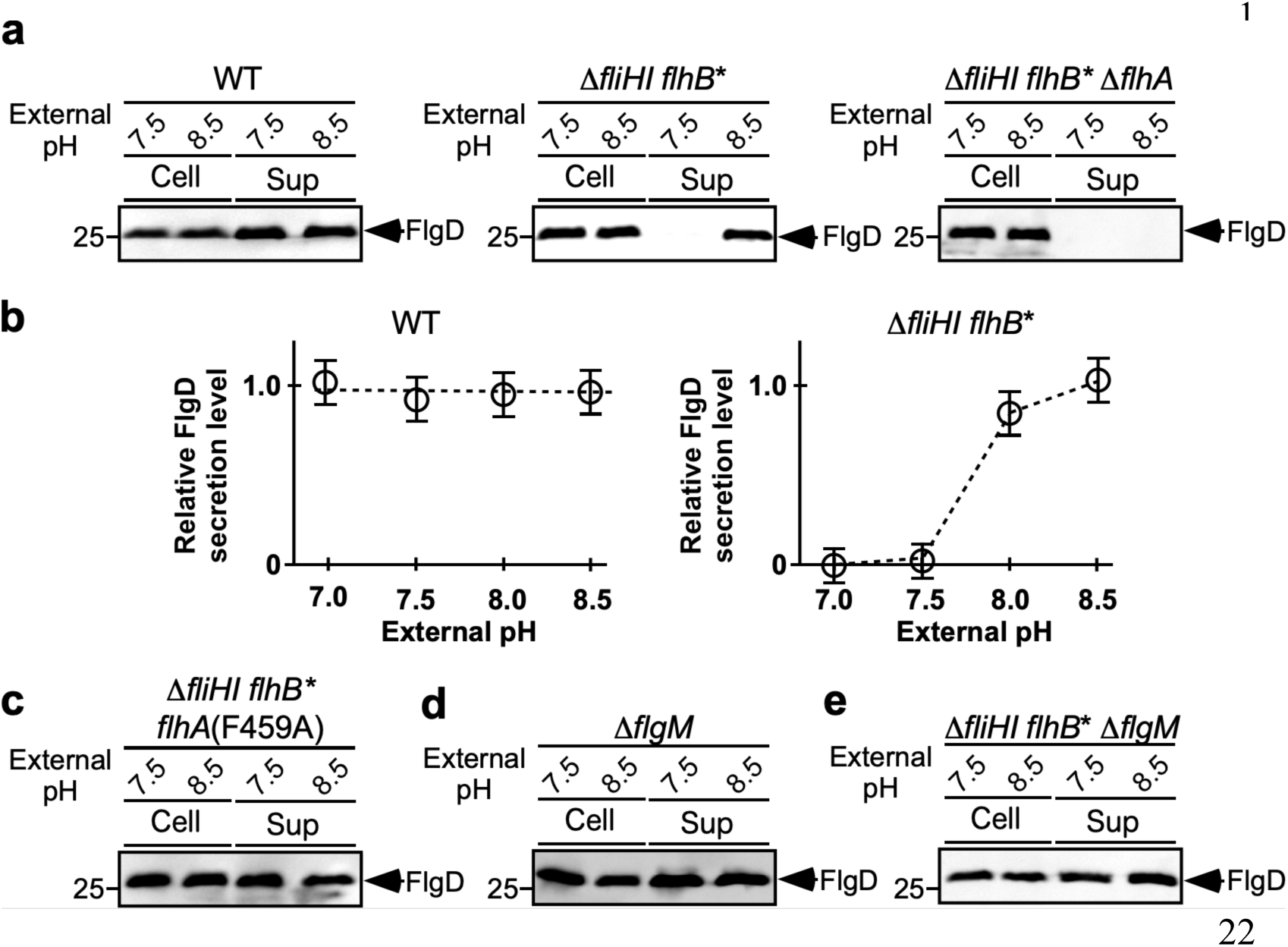
Effect of increase in ΔΨ on flagellar protein export. **(a)** Effect of external pH on flagellar protein export by the export gate complex in the presence and absence of FliH and FliI. Immunoblotting, using polyclonal anti-FlgD antibody, of whole cell proteins (Cell) and culture supernatant fractions (Sup) prepared from SJW1103, MMHI0117 and NH004 (Δ*fliHI flhB*\* Δ*flhA*) grown in TB-7.5 or TB-8.5. **(b)** Relative FlgD secretion levels. FlgD band densities are normalized for the cellular FlgD levels. These data are the average of three independent experiments. **(c, d, f)** Immunoblotting, using polyclonal anti-FlgD antibody, of whole cell proteins (Cell) and culture supernatant fractions (Sup) prepared from **(c)** MMHI0117-1 [Δ*fliHI flhB*\* *flhA(F459A)*], **(d)** MM1103gM (Δ*flgM*) and **(f)** MMHI0117gM (Δ*fliHI flhB*\* Δ*flgM*) grown in TB-7.5 or TB-8.5.

To count the population of flagellated cells and measure their flagellar length in the Δ*fliH-fliI flhB(P28T)* mutant exponentially grown in TB-7.5 or TB-8.5, we labelled flagellar filaments with a fluorescent dye (Fig. 4a) and measured the number and length of the filaments (Supplementary Table 2). Only 1.0% of the Δf*liH-fliI flhB(P28T)* cells had a single flagellar filament at external pH 7.5 (n = 198) (Fig. 4b). In contrast, at external pH 8.5, 60.5% of the Δf*liH-fliI flhB(P28T)* cells produced the filaments with an average number of 1.3 ± 0.5 per cell [mean ± standard deviation (SD), n = 107] (Fig. 4b). The average filament length was 4.8 ± 1.5 μm (n = 50), which was about 2-fold shorter than the wild-type length [9.2 ± 2.2 μm (n = 50)] (Fig. 4c), indicating that the filament growth rate of the Δf*liH-fliI flhB(P28T)* mutant is about 2-fold slower than that of wild-type cells when flagellar construction occurs.

**Fig. 4.**
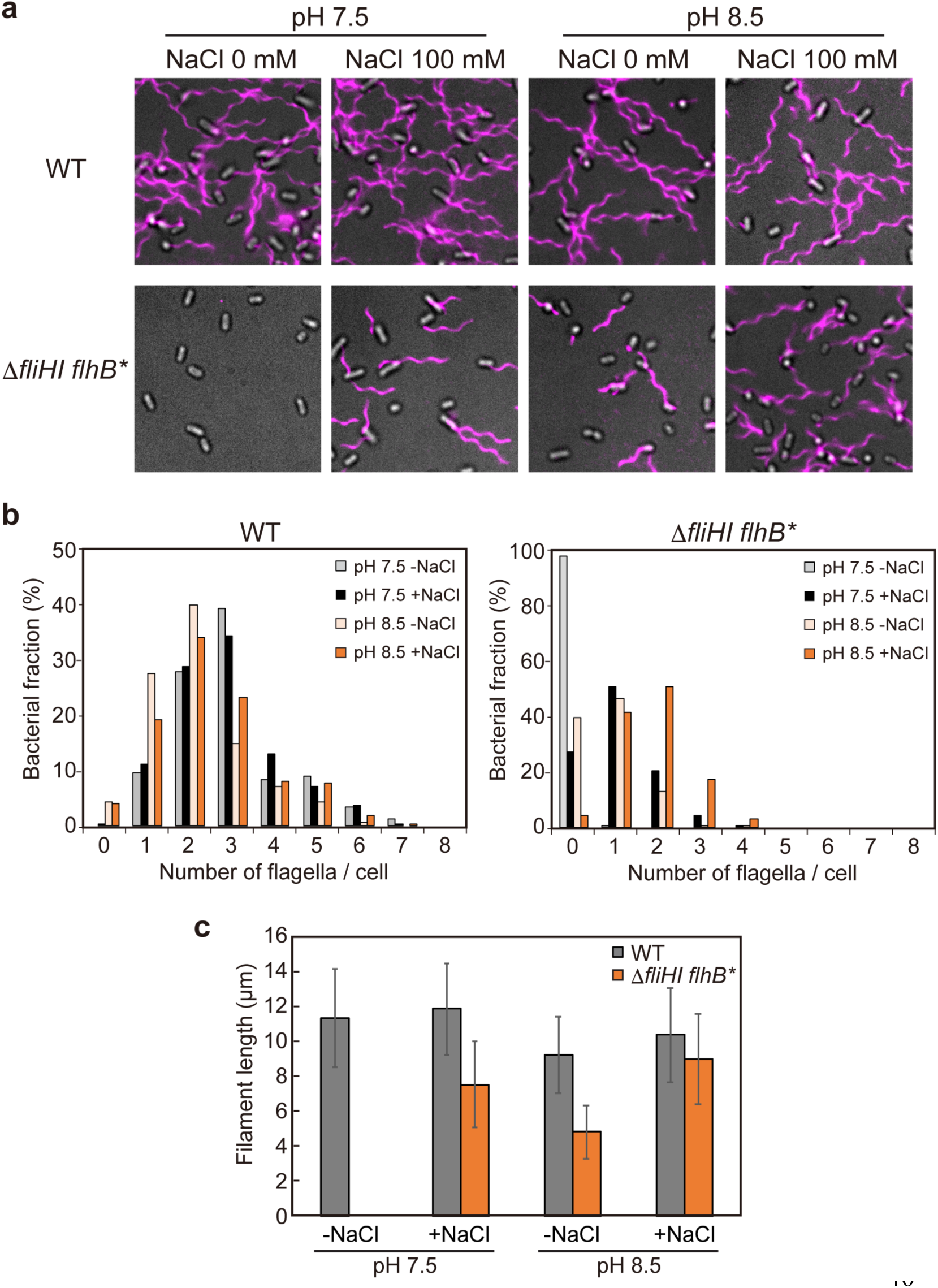
Effect of the greater ΔΨ on flagellar formation. **(a)** Fluorescent images of SJW1103 (WT) and MMHI0117 (Δ*fliHI flhB*\*) grown in TB-7.5 or TB-8.5 with or without 100 mM NaCl. Flagellar filaments were labelled with Alexa Fluor 594. The fluorescence images of the filaments labelled with Alexa Fluor 594 (magenta) were merged with the bright field images of the cell bodies. **(b)** Distribution of the number of the flagellar filaments in the SJW1103 and MMHI0117 cells. More than 150 cells for each strain were counted. **(c)** Measurements of the length of the flagellar filaments. Filament length is the average of 50 filaments, and vertical lines are standard deviations. (See Supplementary Table 2)

The ΔΨ component of PMF is essential for flagellar protein export by *Salmonella* wild-type cells^13,14^. Therefore, we tested whether the 1.5-fold greater ΔΨ also increases the secretion level of FlgD by wild-type cells. Total PMF did not change at all when the external pH increased from 7.5 to 8.5 in the wild-type cells as well (Fig. 2b). An upward pH shift from 7.5 to 8.5 neither affected the secretion of FlgD nor flagellar filament formation significantly (Figs. 3a, b and 4), indicating that the greater ΔΨ does not facilitate protein export by the wild-type protein export apparatus. This suggests that the rate of H^+^-coupled protein translocation by the export gate complex could be high enough to obscure the effect of greater ΔΨ in the presence of FliH and FliI.

### Effect of gain-of-function mutations in FlhA on ΔΨ-dependent flagellar protein export by the Δ*fliH-fliI flhB(P28T)* strain

The Δ*fliH-fliI flhB(P28T)* cells absolutely require the ΔpNa component to drive flagellar protein export at external pH 7.5 because the ΔpH component of PMF is gone^17^. It has been reported that the *flhA(D456V), flhA(F459A)* and *flhA(T490A)* mutations, for which the mutated residues are located in the C-terminal cytoplasmic domain of FlhA (FlhA_C_) (Supplementary Fig. 1a), significantly increase the probability of docking of flagellar building blocks to the export gate complex in the absence of FliH and FliI, thereby increasing the probability of hook-basal body assembly in the Δ*fliH-fliI flhB(P28T)* mutant^21^. This raises the possibility that these *flhA* mutations would reduce Na^+^-dependence of flagellar protein export by the Δ*fliH-fliI flhB(P28T)* mutant. To examine this possibility, we analyzed the effect of Na^+^ depletion on FlgD secretion by the Δ*fliH-fliI flhB(P28T) flhA(F459A)* mutant. This mutant was exponentially grown in TB-7.5 with or without 100 mM NaCl, and then FlgD secretion levels were analyzed by immunoblotting with polyclonal anti-FlgD antibody. FlgD was detected in the culture supernatant in the absence of Na^+^, but the secretion level of FlgD was significantly increased by adding 100 mM NaCl (Supplementary Fig. 1b). We also obtained essentially the same results with alternative Δ*fliH-fliI flhB(P28T) flhA(D456V)* and Δ*fliH-fliI flhB(P28T) flhA(T490M)* mutants (Supplementary Fig. 1b). An upward pH shift from 7.5 to 8.5 to increase ΔΨ did not increase the secretion level of FlgD by the Δ*fliH-fliI flhB(P28T) flhA(F459A)* mutant (Fig. 3c). These results suggest that these *flhA* mutations reduce the ΔΨ-dependency of flagellar protein export by the export gate complex in the absence of FliH and FliI.

We next measured the number and length of the filaments produced by the Δ*fliH-fliI flhB(P28T) flhA(F459A)* mutant grown exponentially in TB-7.5 with or without 100 mM NaCl (Supplementary Fig. 2a and Supplementary Table 3). In the absence of NaCl, 21.8% of the Δ*fliH-fliI flhB(P28T) flhA(F459A)* cells produced the filaments with an average number of 1.1 ± 0.3 per cell (n = 121) while the remaining 78.2% had no filaments (Supplementary Fig. 2b). The average filament length was 3.3 ± 1.6 μm (n = 50) (Supplementary Fig. 2c). In the presence of 100 mM NaCl, about 76.3% of the Δ*fliH-fliI flhB(P28T) flhA(F459A)* mutant cells produced the filaments with an average number of 1.7 ± 0.8 per cell (n = 222) (Supplementary Fig. 2b), indicating that the ΔpNa component of SMF significantly increases the probability of flagellar formation. The average filament length was 3.4 ± 2.0 μm, which was almost the same as that in the absence of NaCl (Supplementary Fig. 2c). We also obtained the same results with the Δ*fliH-fliI flhB(P28T) flhA(D456V)* and Δ*fliH-fliI flhB(P28T) flhA(T490M)* mutants (Supplementary Fig. 2).

To test whether the *flhB(P28T)* and *flhA(F459A)* mutations affect the ΔΨ-dependency of flagellar protein export by the export gate complex in the presence of FliH and FliI, we analyzed the number and length of flagellar filaments produced by the *flhB(P28T)* and *flhA(F459A)* mutants. More than 95% of the *flhB(P28T)* and *flhA(F459A)* mutant cells produced the filaments with the number per cell at the wild-type level in the presence and absence of 100 mM NaCl (Supplementary Fig. 2b). The average filament length of the *flhB(P28T)* mutant was almost the same as that of wild-type cells, but that of the *flhA*(F459A) mutant was about half of the wild-type (Supplementary Fig. 2c) due to the reduced secretion levels of flagellin (FliC) molecules^22^. These observations suggest that the export gate complex does not require either greater ΔΨ or ΔpNa for flagellar protein export and assembly in the presence of FliH and FliI.

Freely diffusing FlhA molecules conduct both H^+^ and Na^+^, but their H^+^ channel activity is lower than the Na^+^ channel activity^17^. To test whether the gain-of-function mutations in FlhA facilitate the H^+^ channel activity of FlhA, we expressed a ratiometric pH indicator probe, pHluorin^23,24^, in *E. coli* BL21 (DE3) cells and measured intracellular pH change upon lowering the external pH value from 7.5 to 5.5 to generate much greater ΔpH (Supplementary Fig. 3a and Supplementary Table 4). Intracellular pH of the FlhA-expressing cells was 7.10 ± 0.06 (mean ± SD), which was ca. 0.06 pH unit lower than that of the vector control (7.16 ± 0.06). This small pH drop was a statistically significant value (*P* = 0.037), indicating the H^+^ channel activity. The Intracellular pH of the cells expressing FlhA with *flhA(D456V), flhA(F459A)* or *flhA(T490A)* mutations were essentially the same as that of the cells expressing wild-type FlhA (Supplementary Fig. 3a), indicating that these *flhA* mutations do not show any increase in the H^+^ channel activity of FlhA.

### Effect of FlgM deletion on flagellar protein export

FlhA_C_ and the C-terminal cytoplasmic domain of FlhB (FlhB_C_) form a docking platform for FliH, FliI, FliJ, export chaperones (FlgN, FliS, FliT) and export substrates (Fig. 1)^25^, and this FlhA_C_-FlhB_C_ docking platform plays an important role in the coordinated flagellar protein export with assembly^26^. We found that the *flhA(D456V), flhA(F459A)* and *flhA(T490A)* mutations in FlhA_C_ activated the export gate complex of the Δ*fliH-fliI flhB(P28T)* mutant to a considerable degree in the absence of greater ΔΨ and ΔpNa (Fig. 3c and Supplementary Fig. 1), raising a question of whether an increase in the expression levels of export substrates also reduces the ΔΨ-dependence of flagellar protein export by the Δ*fliH-fliI flhB(P28T)* mutant. Because depletion of FlgM, which is a negative regulator of the flagellar regulon^27^, considerably increases the expression levels of FliJ, flagellar export chaperones and export substrates, thereby increasing the probability of flagellar formation even in the absence of FliH and FliI^28,29^, we introduced the Δ*flgM*::*km* allele into the wild-type and Δ*fliH-fliI flhB(P28T)* mutant strains by P22-mediated transduction to generate the Δ*flgM* and Δ*fliH-fliI flhB(P28T)* Δ*flgM* mutant strains (Fig. 3d, e). FlgM deletion allowed the Δ*fliH-fliI flhB(P28T)* mutant to secret FlgD into the culture media at external pH 7.5 (Fig. 3e). An upward pH shift from 7.5 to 8.5, which increases ΔΨ, increased the secretion level of FlgD but only by about 1.5-fold. These results suggest that the FlgM deletion significantly reduces the ΔΨ-dependency of ion-coupled protein export by the export gate complex. The greater ΔΨ did not increase the FlgD secretion by the Δ*flgM* mutant in a way similar to wild-type cells (Fig. 3d). Because ΔΨ is essential for flagellar protein export by wild-type cells^13,14^, we suggest that the cytoplasmic ATPase complex consisting of FliH and FliI facilitates efficient and rapid docking of export substrates to the FlhA_C_-FlhB_C_ docking platform in a ΔΨ-dependent manner.

### Effect of greater SMF on flagellar protein export

Flagellar protein export by the Δ*fliH-fliI flhB(P28T)* mutant shows a clear dependence on external Na^+^ concentration at external pH 7.5^17^. Therefore, we tested whether greater SMF increases the secretion level of FlgD by this mutant. Total SMF across the cell membrane was about 45 mV greater at external pH 8.5 than at external pH 7.5 (Fig. 5a and Supplementary Table 1). The Δ*fliH-fliI flhB(P28T)* cells were grown in TB-7.5 at 30°C for 3 hours. After washing twice with TB-7.5, the cells were resuspended in fresh TB-7.5 or TB-8.5 with or without 100 mM NaCl, and incubation was continued at 30°C for 1 hour. In agreement with a previous report^17^, addition of 100 mM NaCl increased the secretion level of FlgD by the Δ*fliH-fliI flhB(P28T)* mutant from almost zero to more than 100-fold at external pH 7.5, and this FlgD secretion level was about 3-fold higher compared to that at external pH 8.5 in the absence of Na^+^ (Fig. 5b). Greater SFM by an upward shift of external pH from 7.5 to 8.5 increased the secretion level of FlgD by about 2-fold in the presence of Na^+^ (Fig. 5b). These results demonstrate that the translocation rate of flagellar building blocks depends on the rate of the Na^+^ flow through the FlhA channel of the export gate complex.

**Fig. 5.**
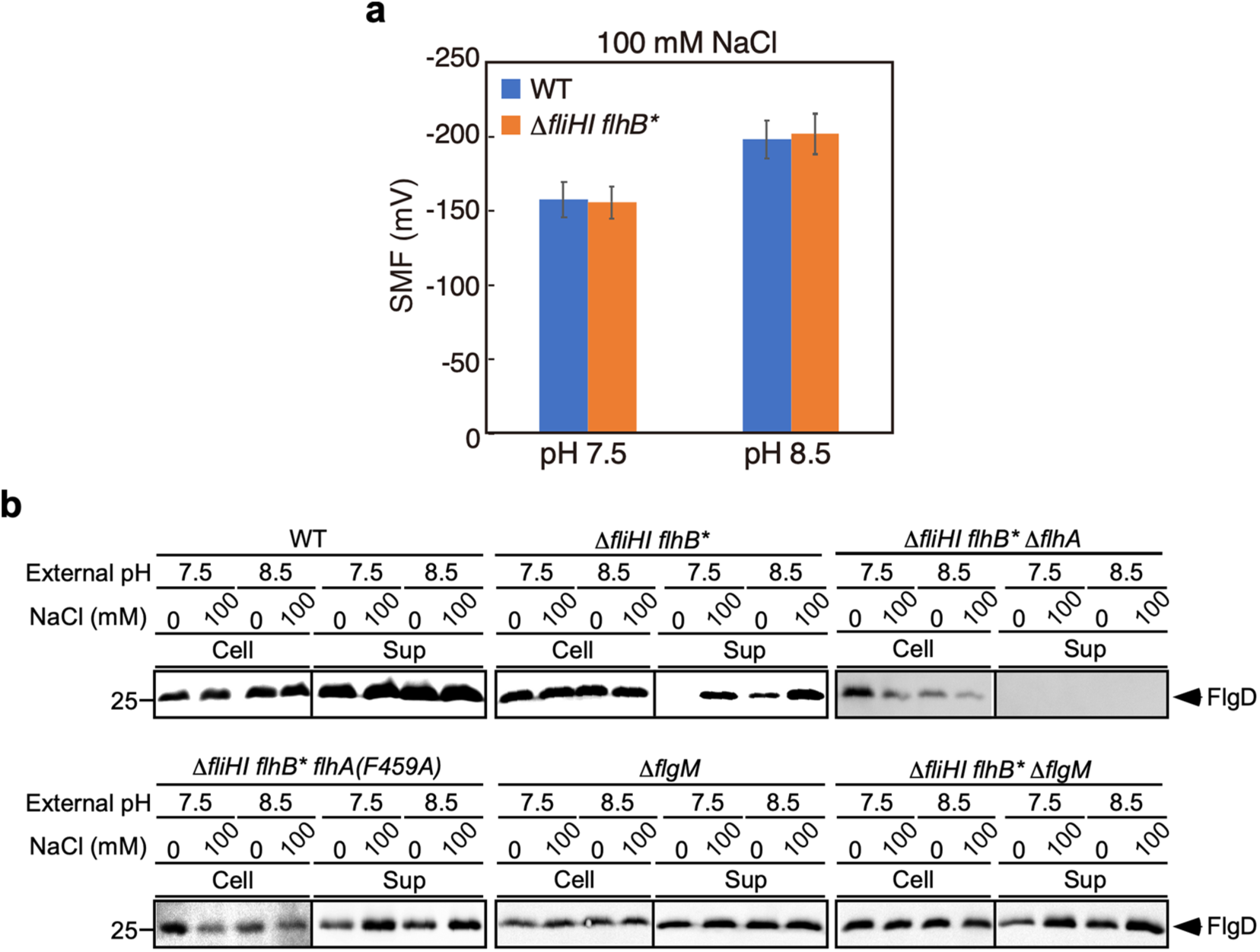
Effect of ΔpNa on flagellar protein export. **(a)** Measurements of SMF. Intracellular Na^+^ concentration was measured with CoroNa green. More than 100 cells were measured. Vertical bars indicate standard deviations. (See Supplementary Table 1) **(b)** Effect of Na^+^ on flagellar protein export at external pH values of 7.5 and 8.5. The SJW1103 (WT), MMHI0117 (Δ*fliHI flhB*\*), NH004 (Δ*fliHI flhB*\* Δ*flhA*), MMHI0117-1 [Δ*fliHI flhB*\* *flhA(F459A)*], MM1103gM (Δ*flgM*) and MMHI0117gM (Δ*fliHI flhB*\* Δ*flgM*) cells were exponentially grown at 30°C in TB-7.5 without 100 mM NaCl. After washing twice with TB-7.5 without 100 mM NaCl, the cells were resuspended in TB-7.5 or TB-8.5 with or without 100 mM NaCl and incubated at 30°C for 1 hour. The whole cell (Cells) and culture supernatant fractions (Sup) were analyzed by immunoblotting with polyclonal anti-FlgD antibody.

We next analyzed the number and length of the flagellar filaments produced by the Δ*fliH-fliI flhB(P28T)* strain in the presence of 100 mM NaCl (Fig. 4a and Supplementary Table 2). About 73.5% of the Δ*fliH-fliI flhB(P28T)* cells produced flagellar filaments with an average of 1.4 ± 0.6 per cell (mean ± SD, n = 125) at external pH 7.5 (Fig. 4b, c). The average filament length was 7.6 ± 2.5 μm (n = 50), which was about 64% of the wild-type length [11.9 ± 2.6 μm (n = 50)] (Fig. 4c). In contrast, about 96.3% of the Δ*fliH-fliI flhB(P28T)* cells produced the filaments with an average of 1.8 ± 0.8 per cell (n = 180) at external pH 8.5 (Fig. 4b). The average filament length was 9.0 ± 2.6 μm (n = 50), which was about 86.5% of the wild-type length [10.4 ± 2.7 μm (n = 50)] (Fig. 4c). Because SMF was larger than PMF under our experimental conditions (Figs 2b and 5a and Supplementary Table 1), we propose that the increase in ΔΨ acts on the FlhA channel to further facilitate the inward-directed ion translocation coupled with the outward-directed protein translocation.

We tested whether the *flhA(F459A)* mutation affects Na^+^-coupled flagellar protein export. The greater SFM did not significantly affect the secretion level of FlgD by the Δ*fliH-fliI flhB(P28T) flhA(F459A)* mutant (Fig. 5b). We also obtained essentially the same results with alternative Δ*fliH-fliI flhB(P28T) flhA(D456V)* and Δ*fliH-fliI flhB(P28T) flhA(T490M)* mutants (Supplementary Fig. 1b). These results raise the possibility that these FlhA mutations do not increase the ΔΨ-dependent Na^+^ channel activity of FlhA. To clarify this, we measured intracellular Na^+^ concentration change of FlhA-expressing *E. coli* cells using a fluorescent Na^+^ indicator dye, CoroNa Green (Supplementary Fig. 3b and Supplementary Table 4). Overexpression of FlhA caused a significant increment in the intracellular Na^+^ concentration in the presence of 100 mM NaCl but not in its absence, in agreement with a previous report^17^. The intracellular Na^+^ concentration of the FlhA-expressing cells increased from 4.5 ± 2.1 mM (average ± standard error, n = 30) to 73.1 ± 10.8 mM (n = 30) (Supplementary Fig. 3b). The intracellular Na^+^ concentration of cells over-expressing FlhA with the D456V, F469A or T490M mutation reached 84.1 ± 16.7 mM (n = 30), 94.2 ± 17.7 mM (n = 30) or 75.6 ± 13.7 mM (n = 30), respectively (Supplementary Fig. 3b), indicating that these *flhA* mutations do not significantly affect the Na^+^ channel activity of FlhA. Therefore, we propose that the inward-directed Na^+^ translocation rate depends not only on SMF across the cell membrane but also on the rate of the outward-directed protein translocation through the polypeptide channel of the transmembrane export gate complex.

We finally investigated whether the greater SMF affects the secretion of FlgD by the Δ*fliH-fliI flhB(P28T)* Δ*flgM* mutant. At external pH 7.5, the secretion level of FlgD was considerably increased by adding 100 mM NaCl (Fig. 5b), indicating that Na^+^ facilitates flagellar protein export. Increasing external pH from 7.5 to 8.5 increased the secretion level of FlgD by about 1.5-fold in the presence of 100 mM NaCl (Fig. 5b). Because the 1.5-fold greater ΔΨ increased the secretion level of FlgD also in the absence of NaCl (Fig. 5b), we suggest that the FlgM deletion increases the docking efficiency of export substrate to the export gate complex in the absence of FliH and FliI and that the ΔpNa component determines the rate of the inward-directed Na^+^ flow coupled with the outward-directed protein translocation through the polypeptide channel. Neither greater ΔΨ nor SMF increased the FlgD secretion by the Δ*flgM* mutant in a way similar to wild-type cells (Fig. 5b). Therefore, we suggest that the cytoplasmic ATPase complex facilitates not only export substrate docking to the export gate complex but also facilitates the H^+^ flow through the FlhA channel.

## Discussion

The transmembrane export gate complex is a dual fuel engine that utilizes both H^+^ and Na^+^ to drive flagellar protein export. FlhA can conduct H^+^ and Na^+^ along IMF across the cell membrane^17^. In the wild-type protein export apparatus, ΔpI (ΔpH or ΔpNa) component of IMF is not essential, and only the ΔΨ component is sufficient for flagellar protein export^13^. But since the ΔpH and ΔpNa components become essential in the absence of the cytoplasmic ATPase complex^14,17^, ΔpH and ΔpNa are thought to be required for efficient transit of H^+^ and Na^+^ across the cell membrane, respectively. However, the role of ΔΨ remained a mystery.

In this study, we used the *Salmonella* Δ*fliH-fliI flhB(P28T)* mutant to see the impact of an increment in the ΔΨ component on flagellar protein export in the absence of ΔpH and ΔpNa, under which condition the flagellar protein export machinery of this mutant is inactive. An upward pH shift from 7.5 to 8.5 increased ΔΨ by 1.5-fold to maintain total PMF (Fig. 2). When ΔΨ rose above a certain threshold, the transmembrane export gate complex of the Δ*fliH-fliI flhB(P28T)* mutant became an active protein transporter to drive H^+^-coupled protein export to form a few flagella in the absence of positive ΔpH and ΔpNa (Figs. 3b and 4). This suggests that the export gate complex is a voltage-gated protein transporter to open its polypeptide and ion channels in a ΔΨ-dependent manner. The *flhA(F459A)* mutation in FlhA_C_, which increases the probability of the substrate entry into a polypeptide channel of the export gate complex in the absence of FliH and FliI^21^, allowed the export gate complex of the Δ*fliH-fliI flhB(P28T)* mutant to transport FlgD to the cell exterior to a considerable degree in the absence of ΔpH and ΔpNa (Fig. 2c and Supplementary Fig. 1). Furthermore, a large increase in the expression levels of export substrates by FlgM depletion also increased the secretion level of FlgD by the Δ*fliH-fliI flhB(P28T)* mutant in the absence of ΔpH and ΔpNa (Fig. 2c). These results suggest that the efficient docking of export substrates to the FlhA_C_-FlhB_C_ docking platform autonomously induces the gate opening of the polypeptide channel, thereby facilitating the outward-directed protein translocation through the polypeptide channel coupled with the inward-directed H^+^ translocation through the FlhA channel, but there is also a ΔΨ-dependent activation mechanism of protein export that independently works to compensate other mechanisms to maintain the protein export activity under some conditions. Therefore, we propose that ΔΨ acts not only on the polypeptide channel to drive the translocation of export substrates across the cytoplasmic membrane but also on the FlhA ion channel to facilitate the H^+^ flow coupled with flagellar protein export.

Neither H^+^ nor Na^+^ channel activity of FlhA was affected by the *flhA(F459A)* mutation (Supplementary Fig. 3). However, the ΔpNa component of SMF increased the secretion level of FlgD by the Δ*fliH-fliI flhB(P28T) flhA(F459A)* mutant at both external pH 7.5 and 8.5 (Fig. 5b and Supplementary Fig. 1), suggesting that ΔpNa facilitates the Na^+^ translocation through the FlhA channel coupled with the protein translocation through the polypeptide channel. In the presence of FliH and FliI, the greater ΔΨ above the threshold affected the secretion level of FlgD neither by the *flhA(F450A)* gain-of-function mutation nor the FlgM deletion (Figs 3 and 5b). Therefore, we suggest that the function of the cytoplasmic ATPase complex is sufficient to make the export gate complex fully functional by facilitating the docking of export substrates to the FlhA_C_-FlhB_C_ docking platform and subsequent substrate entry into a polypeptide channel and that the free energy derived from ATP hydrolysis by the FliI ATPase is required for rapid and efficient ΔΨ-dependent H^+^ translocation through the FlhA channel, which is tightly coupled with the translocation of export substrates into a polypeptide channel as proposed previoulsy^12,15,16,21^.

Why does the flagellar export engine maintain the ΔΨ-dependent activation mechanism throughout the evolutionary process? When planktonic motile cells attach to a solid surface, 3’-5’ cyclic diguanylate monophosphate (cyclic-di-GMP), a nucleotide second messenger, inhibits flagella-driven motility to trigger a motility-to-biofilm transition. The flagellar regulon is placed under control of cyclic-di-GMP signalling networks so that flagellar gene transcription is suppressed during biofilm development^30,31^. It has been reported that cyclic di-GMP directly binds to the ATP-binding pocket of the FliI ATPase to inhibit the ATPase activity, suggesting that the FliI ATPase is not functional in cells living in the biofilm^32^. Furthermore, ΔΨ is quite small in the biofilm structure^20^. On the other hand, flagella-driven motility of a subpopulation of planktonic cells in the biofilm structure is essential to keep the cells in the biofilm alive and healthy^33^, raising a question of how the flagellar export engine is activated to generate flagellated cells during biofilm development. It has been shown that a metabolic trigger induces release of intracellular potassium (K^+^) through a K^+^ channel, which in turn depolarizes neighbouring cells in the biofilm, thereby allowing the cells to uptake nutrients in a ΔΨ-dependent manner^20^. Because the FliI ATPase adopts an inactive form and available flagellar building blocks are limited in the cytoplasm of the cells in the biofilm, we propose that the voltage-gated mechanism of the flagellar export engine is essential for survival of the cells by efficiently generating flagellated cells in the biofilm structure.

## Methods

### Bacterial strains, plasmids and Media

*Salmonella* strains and plasmids used in this study are listed in Supplementary Table 4. T-broth (TB) contained 1% Bacto tryptone, 10 mM potassium phosphate, pH 7.5 (TB-7.5) or pH 8.5 (TB-8.5).

### Secretion assay

*Salmonella* cells were grown overnight in TB-7.5 without 100 mM NaCl. A 50 μl of the overnight culture was inoculated into a 5 ml of fresh TB-7.5 or TB-8.5 with or without 100 mM NaCl and incubated at 30 °C with shaking until the cell density had reached an OD_600_ of ca. 1.4–1.6. To test the effect of a chemical gradient of Na^+^ on flagellar protein export, the cells were grown in 5 ml of TB-7.5 containing 100 mM NaCl with shaking at 30 °C until the cell density had reached an OD_600_ of ca. 1.0–1.2. After washing the cells twice, the cells were resuspended in 5 ml TB-7.5 or TB-8.5 with or without 100 mM NaCl, followed by incubation at 30 °C for 1 hour with shaking. Cultures were centrifuged to obtain cell pellets and culture supernatants. Cell pellets were resuspended in an SDS-loading buffer (62.5 mM Tris-HCl, pH 6.8, 2% SDS, 10% glycerol, 0.001% bromophenol blue) containing 1 μl of 2-mercaptoethanol, normalized to a cell density to give a constant number of cells. Proteins in the culture supernatants were precipitated by 10% trichloroacetic acid, suspended in a Tris/SDS loading buffer (one volume of 1 M Tris, nine volumes of 1 X SDS loading buffer) containing 1 μl of 2-mercaptoethanol and heated at 95°C for 3 min. After Sodium Dodecyl Sulfate–polyacrylamide gel electrophoresis (SDS–PAGE), immunoblotting with polyclonal anti-FlgD antibody was carried out as described previously^34^. Detection was performed with an ECL plus immunoblotting detection kit (GE Healthcare).

### Observation of flagellar filaments with a fluorescent dye

The flagellar filaments produced by *Salmonella* cells were labelled using anti-FliC antiserum and anti-rabbit IgG conjugated with Alexa Fluor^®^ 594 (Invitrogen) as described previously^16^. The cells were observed by fluorescence microscopy as described previously^35^. Fluorescence images were analysed using ImageJ software version 1.52 (National Institutes of Health).

### Measurements of ΔΨ, intracellular pH and intracellular sodium ion concentration

The ΔΨ component was measured using tetramethylrhodamine methyl ester (Invitrogen) as described previously^14^. Intracellular pH measurements with a ratiometric fluorescent pH indicator protein, pHluorin(M153R), were carried out as described before ^24^. Intracellular sodium ion concentration was measured using CoroNa Green (Invitrogen) as described previously^17,36^.

### Statistical analysis

Statistical analyses were done using Prism 7.0c software (GraphPad). Comparisons were performed using a two-tailed Student’s *t*-test. A *P* value of < 0.05 was considered to be statistically significant difference.

## Acknowledgements

We acknowledge Yasuyo Abe for technical assistance. This work was supported in part by JSPS KAKENHI Grant Numbers JP26293097 and JP19H03182 (to T.M.), JP15H05593 and JP18K06159 (to Y.V.M), JP18K14638 and JP20K15749 (to M.K.) and JP25000013 (to K.N.) and MEXT KAKENHI Grant Numbers JP15H01640 (to T.M.) and JP26115720 and JP15H01335 (to Y.V.M). This work has also been partially supported by JEOL YOKOGUSHI Research Alliance Laboratories of Osaka University to K.N.

## Author Contributions

T.M. and K.N. conceived and designed research; T.M., Y.V.M. and M.K. performed experiments; T.M., Y.V.M. and M.K. analysed the data, and T.M. and K.N. wrote the paper based on discussion with other authors.

