## Supplementary Information for "Membrane voltage-dependent activation of the flagellar protein export engine"

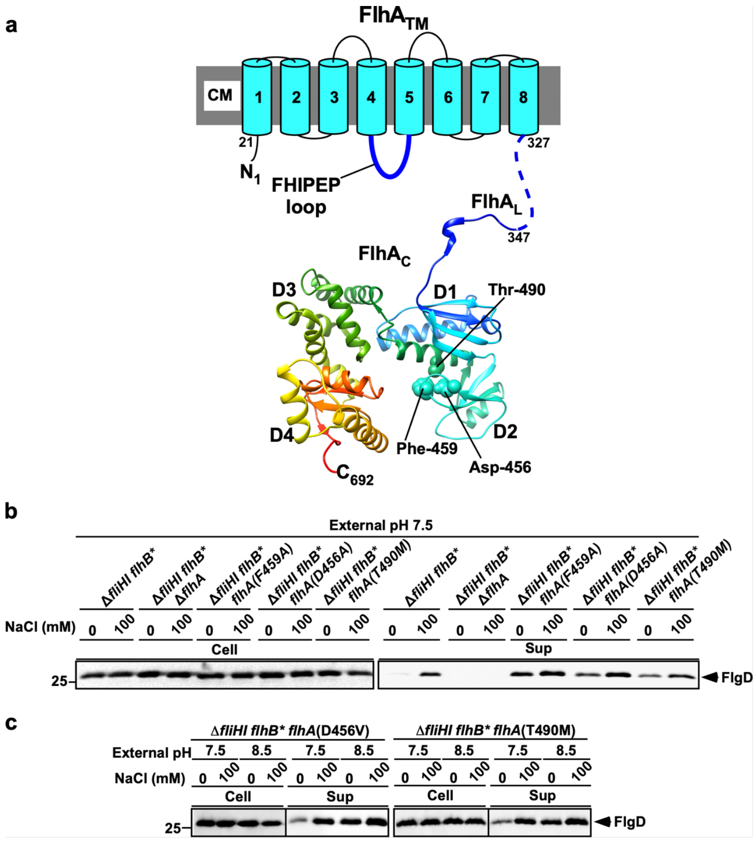

**Supplementary Fig. 1. Effect of gain-of-function mutations in FlhA on flagellar protein export. (a)** FlhA structure. FlhA is composed of an N-terminal transmembrane region ($FlhA_{TM}$) and a large C-terminal cytoplasmic domain ($FlhA_C$). $FlhA_C$ forms a docking platform for FliH, FliI, FliJ, export chaperones and export substrates along with the C-terminal cytoplasmic domain of FlhB ($FlhB_C$). $FlhA_C$ (PDB ID, 3A5I) consists of four domains, D1, D2, D3 and D4 and a flexible linker ($FlhA_L$). The Cα backbone is color-coded from blue to red, going through the rainbow colors from the N- to the C-terminus. Asp-456, Phe-459 and Thr-490 residues, which are well conserved among FlhA homologues, are responsible for the interaction of $FlhA_C$ with flagellar export chaperone–substrate complexes. A highly conserved FHIPEP loop between transmembrane helices 4 and 5 of $FlhA_{TM}$ binds to $FlhA_C$ and $FlhB_C$ to coordinate flagellar protein export with assembly. **(b)** Immunoblotting, using polyclonal anti-FlgD antibody, of whole cell proteins (Cell) and culture supernatant fractions (Sup) prepared from MMHI0117 (Δ*fliHI flhB**), NH004 (Δ*fliHI flhB** Δ*flhA*), MMHI0117-1 [Δ*fliHI flhB* flhA(F459A)*], MMHI0117-2 [Δ*fliHI flhB* flhA(D456V)*] and MMHI0117-3 [Δ*fliHI flhB* flhA(T490M)*] grown exponentially at 30ºC in TB-7.5 with or without 100 mM NaCl. **(c)** Effect of $Na^+$ on flagellar protein export at external pH values of 7.5 and 8.5. The MMHI0117-2 and MMHI0117-3 cells were exponentially grown at 30ºC in TB-7.5 without 100 mM NaCl. After washing twice with TB-7.5 without 100 mM NaCl, the cells were resuspended in TB-7.5 or TB-8.5 with or without 100 mM NaCl and incubated at 30ºC for 1 hour. The whole cell (Cells) and culture supernatant fractions (Sup) were analyzed by immunoblotting with polyclonal anti-FlgD antibody.

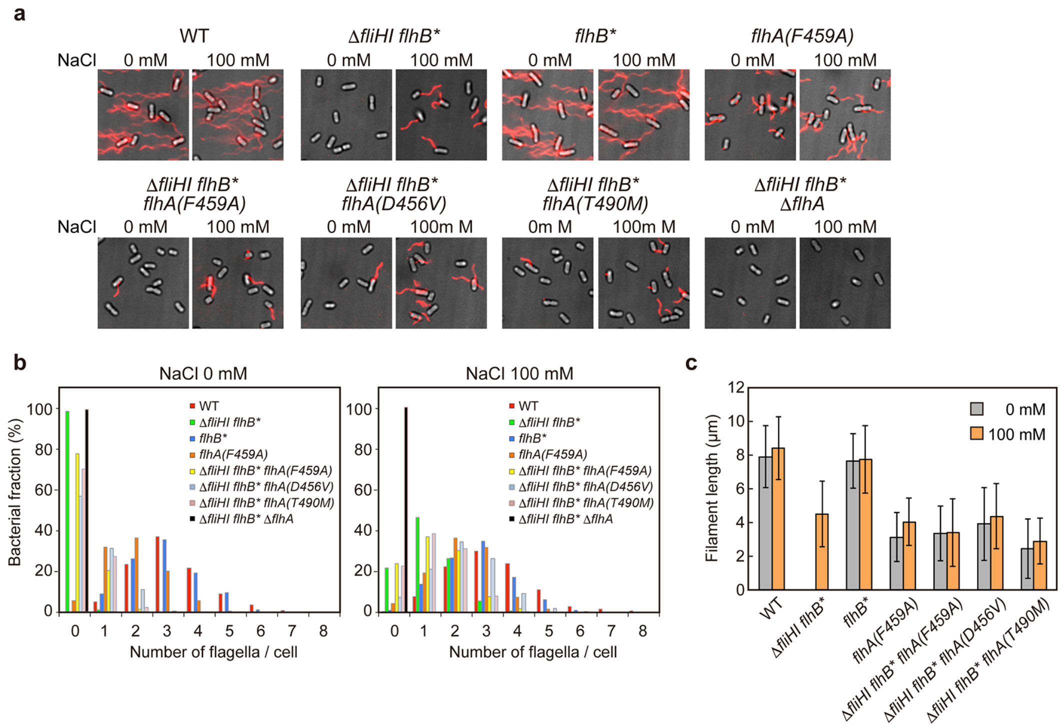

**Supplementary Fig. 2. Effect of gain-of-function mutations in FlhA on flagellar filament formation by the Δ*fliH-fliI flhB(P28T)* strain. (a)** Fluorescent images of SJW1103 (indicated as WT), MMHI0117 (indicated as Δ*fliHI flhB**), MMB017 (indicated as *flhB**), MMA459 [indicated as *flhA(F459A)*], MMHI0117-1 [Δ*fliHI flhB* flhA(F459A)*], MMHI0117-2 [Δ*fliHI flhB* flhA(D456V)*], MMHI0117-3 [Δ*fliHI flhB* flhA(T490M)*] and NH004 (indicated as Δ*fliHI flhB** Δ*flhA*). Flagellar filaments were labelled with Alexa Fluor 594. The fluorescence images of the filaments labelled with Alexa Fluor 594 (red) were merged with the bright field images of the cell bodies. **(b)** Distribution of the number of the flagellar filaments. More than 200 cells for each strain were counted. **(c)** Measurements of the length of the flagellar filaments. Filament length is the average of flagellar filaments for each strain, and vertical lines are standard deviations. (See Supplementary Table 3)

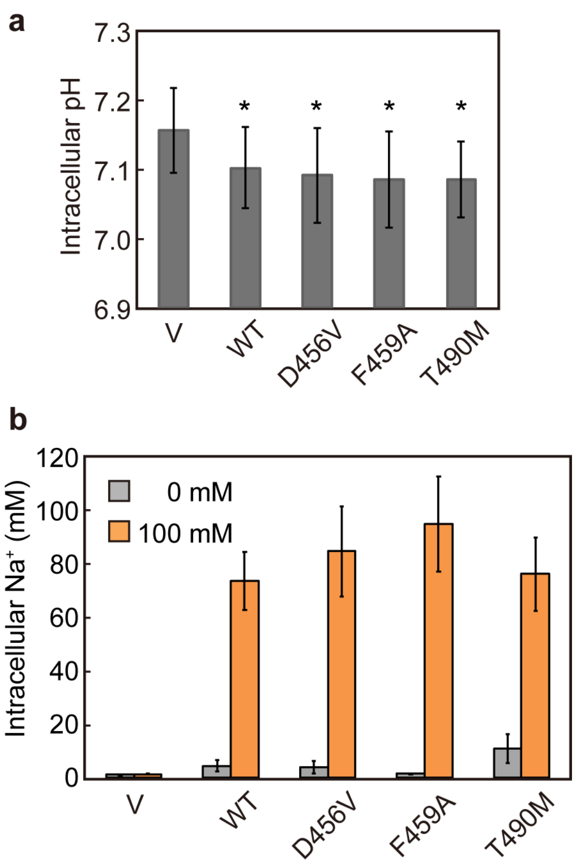

**Supplementary Fig. 3. Effect of gain-of-function mutations in FlhA on the $H^+$ and $Na^+$ channel activities of FlhA. (a)** Effect of overexpression of FlhA on intracellular pH. Intracellular pH was measured with pHluorin at an external pH of 5.5. The BL21(DE3) strain expressing the pHluorin probe was transformed with pBAD24 (Vector, V), pNH319 (FlhA, indicated as WT), pNH319(D456V), [FlhA(D456V), indicated as D456V], pNH319(F459A) [FlhA(F459A), indicated as F459A] or pNH319(T490M) [FlhA(T490M), indicated as T490M]. Vertical bars indicate standard deviations of twelve independent biological replicates. The data that exhibited a statistically significant intracellular change compared with the vector control are highlighted with an asterisk ($P < 0.05$). **(b)** Effect of overexpression of FlhA on intracellular $Na^+$ concentration. Intracellular $Na^+$ concentration was measured with CoroNa Green in the presence and absence of 100 mM NaCl at external pH 7.0. The BL21(DE3) strain was transformed with the above plasmids. For each transformants, 30 cells were measured. Vertical bars indicate standard errors. (See Supplementary Table 4)

**Supplementary Table 1. Ion motive force in *Salmonella* cells**

| | $pH_{ex}$ | 7.5 | | 8.5 | |
|---|---|---|---|---|---|
| | $[Na^+]_{ex}$ (mM) | 0 | 100 | 0 | 100 |
| SJW1103 (WT) | $pH_{in}$ | 7.41 ± 0.05 (n = 4) | 7.41 ± 0.05 (n = 4) | 7.46 ± 0.13 (n = 4) | 7.46 ± 0.13 (n = 4) |
| | Membrane potential (mV) | -91.7 ± 5.0 (n = 165) | -93.5 ± 4.0 (n = 215) | -135.2 ± 4.9 (n = 191) | -134.3 ± 5.3 (n = 56) |
| | $[Na^+]_{in}$ (mM) | 4.2 ± 0.4 | 8.0 ± 12.9 | 4.2 ± 0.4 | 8.0 ± 12.9 |
| | PMF (mV) | -86.5 ± 7.9 | -88.3 ± 6.8 | -74.2 ± 12.3 | -73.2 ± 12.7 |
| | SMF (mV) | - | -157.8 ± 11.5 | - | -198.5 ± 12.8 |
| MMHI0117 [Δ*fliH-fliI flhB(P28T)*] | $pH_{in}$ | 7.43 ± 0.12 (n = 4) | 7.43 ± 0.12 (n = 4 ) | 7.54 ± 0.03 (n = 4) | 7.54 ± 0.03 (n = 4) |
| | Membrane potential (mV) | -92.1 ± 5.6 (n = 71) | -91.6 ± 3.3 (n = 270) | -140.7 ± 5.0 (n = 201) | -137.7 ± 6.0 (n = 144) |
| | $[Na^+]_{in}$ (mM) | 4.2 ± 0.4 | 8.0 ± 12.9 | 4.2 ± 0.4 | 8.0 ± 12.9 |
| | PMF (mV) | -88.1 ± 10.5 | -87.6 ± 10.5 | -84.5 ± 4.8 | -81.5 ± 7.5 |
| | SMF (mV) | - | -155.8 ± 10.9 | - | -201.9 ± 13.6 |

Ion motive force is defined as equation [1].

$$IMF = V_m + \frac{k_B T}{q} \ln \frac{[ion]_{in}}{[ion]_{ex}} \qquad [1]$$

where $V_m$ and $q$ are $\Delta\psi$ and charge of ions, respectively. $[ion]_{in}$ and $[ion]_{out}$ are the internal and external ion concentrations, respectively.

**Supplementary Table 2. Effects of external pH and external $Na^{+}$ concentration on flagellar formation.**

| | $pH_{ex}$ | NaCl (mM) | Fraction of flagellated cells (%) | Average number of flagella in flagellated cell (mean ± SD) | Average length of filament (μm) (mean ± SD) |
|---|---|---|---|---|---|
| SJW1103 (WT) | 7.5 | 0 | 100 (n = 164) | 2.9 ± 1.2 (n = 164) | 11.3 ± 2.8 (n = 50) |
| | | 100 | 100 (n = 177) | 2.9 ± 1.3 (n = 177) | 11.9 ± 2.6 (n = 50) |
| | 8.5 | 0 | 100 (n = 152) | 2.1 ± 1.2 (n = 152) | 9.2 ± 2.2 (n = 50) |
| | | 100 | 100 (n = 192) | 2.5 ± 1.4 (n = 192) | 10.4 ± 2.7 (n = 50) |
| MMHI0117 (Δ*fliHI flhB**) | 7.5 | 0 | 1.0 (n = 198) | 1.0 ± 0.0 (n = 2) | - |
| | | 100 | 73.5 (n = 170) | 1.4 ± 0.6 (n = 125) | 7.6 ± 2.5 (n = 50) |
| | 8.5 | 0 | 60.5 (n = 177) | 1.3 ± 0.5 (n = 107) | 4.8 ± 1.5 (n = 50) |
| | | 100 | 96.3 (n = 187) | 1.8 ± 0.8 (n =1 80) | 9.0 ± 2.6 (n = 50) |

$pH_{ex}$, external pH

**Supplementary Table 3. Effect of $Na^+$ ions on average number and length of flagellar filaments**

| | NaCl (mM) | Fraction of flagellated cells (%) | Average number of flagella in flagellated cell (± SD) | Average length of filament (µm ± SD) |
|---|---|---|---|---|
| SJW1103 (WT) | 0 | 100 (n = 320) | 3.2 ± 1.2 (n = 320) | 7.8 ± 1.8 (n = 50) |
| | 100 | 99.6 (n = 264) | 3.3 ± 1.3 (n = 263) | 8.4 ± 1.9 (n = 50) |
| MMHI0117 (Δ*fliHI flhB**) | 0 | 1.1 (n = 371) | 1.0 ± 0.4 (n = 4) | - |
| | 100 | 78.4 (n = 338) | 1.5 ± 0.7 (n = 265) | 4.5 ± 1.9 (n = 50) |
| MMB017 (*flhB**) | 0 | 100 (n = 214) | 3.0 ± 1.1 (n = 214) | 7.6 ± 1.6 (n = 110) |
| | 100 | 99.1 (n = 227) | 2.8 ± 1.1 (n = 225) | 7.7 ± 2.0 (n = 110) |
| MMA459 [*flhA(F459A)*] | 0 | 94.5 (n = 253) | 2.0 ± 0.9 (n = 239) | 3.1 ± 1.5 (n = 110) |
| | 100 | 95.7 (n = 208) | 2.3 ± 0.9 (n = 199) | 4.0 ± 1.4 (n = 110) |
| MMHI0117-1 [Δ*fliHI flhB* flhA(F459A)*] | 0 | 21.8 (n = 555) | 1.1 ± 0.3 (n = 121) | 3.3 ± 1.6 (n = 18) |
| | 100 | 76.3 (n = 291) | 1.7 ± 0.8 (n = 222) | 3.4 ± 2.0 (n = 58) |
| MMHI0117-2 [Δ*fliHI flhB* flhA(D456V)*] | 0 | 42.9 (n = 515) | 1.3 ± 0.5 (n = 221) | 3.9 ± 2.1 (n = 38) |
| | 100 | 92.6 (n = 444) | 2.4 ± 1.8 (n = 411) | 4.4 ± 1.9 (n = 52) |
| MMHI0117-1 [Δ*fliHI flhB* flhA(T490M)*] | 0 | 29.4 (n = 459) | 1.1 ± 0.3 (n = 135) | 2.4 ± 1.7 (n = 47) |
| | 100 | 77.4 (n = 526) | 1.6 ± 0.7 (n = 407) | 2.9 ± 1.4 (n = 50) |
| NH004 (Δ*fliHI flhB** Δ*flhA*) | 0 | 0 (n = 356) | - | - |
| | 100 | 0 (n = 238) | - | - |

**Supplementary Table 4. Ion conductivity of FlhA mutants.**

| | | V | WT | D456V | F459A | T490M |
|---|---|---|---|---|---|---|
| Internal pH (mean ± SD) | | 7.16 ± 0.06 (n = 12) | 7.10 ± 0.06 (n = 12) | 7.09 ± 0.07 (n = 12) | 7.09 ± 0.07 (n = 12) | 7.09 ± 0.05 (n = 12) |
| $[Na^+]_{in}$ (mM) (mean ± SEM) | NaCl, 0 mM | 1.48 ± 0.03 (n = 30) | 4.5 ± 2.1 (n=30) | 4.0 ± 2.3 (n = 30) | 1.6 ± 0.1 (n = 30) | 10.9 ± 5.3 (n = 30) |
| | NaCl, 100 mM | 1.51 ± 0.03 (n = 30) | 73.1 ± 10.8 (n = 30) | 84.1 ± 16.7 (n = 30) | 94.2 ± 17.7 (n = 30) | 75.6 ± 13.7 (n = 30) |

V, vector control.

### Supplementary Table 5. Strains and plasmids used in this study

| Strains/ Plasmids | Relevant characteristics | Source or reference |
|---|---|---|
| ***E. coli*** | | |
| BL21 (DE3) | Over-expression of proteins | Novagen |
| ***Salmonella*** | | |
| SJW1103 | Wild type for motility and chemotaxis | 1 |
| MM1103gM | SJW1103 Δ*flgM*::*Km* | This study |
| MMA459 | *flhA(F459A)* | 2 |
| MMB017 | *flhB(P28T)* | 3 |
| MMHI0117 | Δ*fliH-fliI flhB(P28T)* | 3 |
| MMHI0117-1 | Δ*fliH-fliI flhB(P28T) flhA(F459A)* | 2 |
| MMHI0117-2 | Δ*fliH-fliI flhB(P28T) flhA(D456V)* | 2 |
| MMHI0117-3 | Δ*fliH-fliI flhB(P28T) flhA(T490M)* | 2 |
| MMHI0117gM | Δ*fliH-fliI flhB(P28T)* Δ*flgM*::*Km* | This study |
| NH004 | Δ*fliH-fliI flhB(P28T)* Δ*flhA* | 4 |
| **Plasmids** | | |
| pBAD24 | Expression vector | 5 |
| pNH319 | pBAD24/ N-His-FLAG-FlhA | 6 |
| pNH319(D456V) | pBAD24/ N-His-FLAG-FlhA(D456V) | This study |
| pNH319(F459A) | pBAD24/ N-His-FLAG-FlhA(F459A) | This study |
| pNH319(T490M) | pBAD24/ N-His-FLAG-FlhA(T490M) | This study |
| pYC17 | pACTrc/pHluorin | 6 |
| pYVM001 | pKK223-3/pHluorin(M153R) | 7 |
